# Cellular modifiers of TDP-43 phase transition and cytoplasmic aggregation

**DOI:** 10.64898/2025.12.16.694670

**Authors:** Natalie Chin, Qi Zhang, Jizhong Zou, Ken Chih-Chien Cheng, Wei Zheng, Yihong Ye

## Abstract

RNA-binding protein TAR DNA-binding protein 43 (TDP-43) can form liquid-like, nuclear assemblies whose phase behavior is thought to influence its aggregation propensity and neurotoxic activity. The mechanism(s) that governs the liquid-to-solid phase transition of TDP-43 is poorly defined. Here we combined chemical and genome-wide genetic screenings to identify cellular factors that modulate the phase behavior of an RNA-binding defective TDP-43 mutant. Our screens uncovered multiple cellular processes including RNA splicing, protein translation, proteostasis imbalance and nuclear export as TDP-43 phase regulators. We also developed a semi-permeabilized cell system that partially recapitulates the TDP-43 phase transition in vitro, and showed that nuclear export inhibition reshapes the nuclear environment to favor RNA-dependent liquid-liquid phase separation (LLPS) of TDP-43, which mitigates its aggregation. Nuclear export inhibition in a brain organoid model bearing an ALS-associated mutation reduces pathogenic phospho-TDP-43 accumulation. These findings identify multiple modulators of TDP-43 phase transitions in a sensitized model system and establish a framework for further dissecting the link between nuclear transport and TDP-43 phase dynamics.

---

The RNA-binding protein TAR DNA-binding protein 43 (TDP-43), encoded by the *TARDBP* gene, plays a central role in RNA transport, splicing and metabolism (Guo and Shorter, 2017; Suk and Rousseaux, 2020), and is the major pathological component of cytoplasmic inclusions in amyotrophic lateral sclerosis (ALS) and frontotemporal dementia (FTD) (Kellett et al., 2025; Neumann et al., 2006; Rummens and Da Cruz, 2025). TDP-43 contains two RNA-recognition motifs (RRMs), a bipartite nuclear-localization sequence, and a glycine-rich C-terminal low-complexity domain (LCD) that is intrinsically prone to aggregation (Tziortzouda et al., 2021). Under physiological conditions, TDP-43 binds RNAs in the nucleus, participating in multiple RNA-metabolic processes. In contrast, disease-causing mutations trigger its mis-localization to the cytoplasm, forming detergent-resistant inclusions. TDP-43-positive protein aggregates were present in ∼97% of ALS and ∼45% of FTD cases, as well as in a subset of Alzheimer’s disease patients. These diseases are now collectively termed TDP-43 proteinopathies (Brettschneider et al., 2014; Neumann et al., 2006). Both loss of nuclear function and gain of cytoplasmic toxicity have been implicated in neurodegeneration (Suk and Rousseaux, 2020; Tziortzouda et al., 2021), underscoring the importance of nucleocytoplasmic transport in regulating TDP-43 patho-physiological functions.

Like many RNA-binding proteins (RBPs), TDP-43 undergoes liquid-liquid phase separation to form membraneless biomolecular condensates (Conicella et al., 2016; Shorter, 2019; Tziortzouda et al., 2021). Under conditions of cellular stress or in the presence of RNA-binding mutations, these condensates can transition from dynamic liquid droplets to rigid gel-like or solid assemblies (Conicella et al., 2016; Gasset-Rosa et al., 2019; Lang et al., 2024). To date, more than 70 disease-linked mutations in *TARDBP* have been identified in ALS or FTD, many within LCD, while others clustering near or in RRMs to disrupt RNA binding (Tziortzouda et al., 2021). Defects in RNA binding appears to be a major determinant of TDP-43 phase behavior (Babinchak et al., 2019; Mann et al., 2019). Disease-associated mutations diminish TDP-43’s activities in RNA metabolism, causing widespread cryptic exon inclusion (Ling et al., 2015; Ma et al., 2022; Mehta et al., 2023; Zhang et al., 2025b), alternative polyadenylation and other RNA maturation defects that collectively contribute to neurotoxicity (Bryce-Smith et al., 2025; Zeng et al., 2025).

Apart from disease-associated mutations, post-translational modifications (PTM), especially acetylation, have been shown to modulate TDP-43’s RNA binding capacity (Cohen et al., 2015; Zhang et al., 2025b). Acetylation was detected on endogenous TDP-43 in ALS patient samples. Although the abundance of acetylated TDP-43 has not been determined, acetylated TDP-43 is enhanced in cells exposed to oxidative or proteotoxic stress (Cohen et al., 2015; Wang et al., 2017). Interestingly, like disease-associated TDP-43 mutants, acetylated TDP-43 is also more aggregation-prone and displays a similar phase regulation pattern (Cohen et al., 2015). Moreover, a recent study showed that a TDP-43 acetylation mimetic mutant designated as 2KQ behaves similarly as RNA-binding defective TDP-43 variants as they all undergo a unique form of demixing, forming “anisosomes” that contain an anisotropic spherical shell of TDP-43 and a central liquid core enriched in the HSP70 chaperone (Yu et al., 2021).

Anisosomes were observed mostly in the nucleus under overexpression conditions. Although their existence in humans and relevance to ALS or FTD are unclear, TDP43 2KQ offers a platform for dissecting the potential link between TDP-43 phase behaviors (droplet formation, gel structures etc.) and aggregation (Yu et al., 2021).

In this study, we combined a chemical-genetic screen approach with a genome-wide siRNA screen to identify molecules that modulate the phase behavior of TDP-43 2KQ. Our study reveals critical contributions of RNA splicing, protein translation, and the HSP90/ubiquitin–proteasome proteostasis network to TDP-43 phase transition. Additionally, nuclear export inhibition can influence the transition of TDP-43 from a nuclear liquid-demixed form to a cytoplasm-localized immobile gel-like structure akin to protein aggregation. In an iPSC-derived organoid model bearing an ALS-associated mutation (Zhang et al., 2025b), nuclear export inhibition reduces cytoplasmic phosphorylated-TDP-43. Collectively, these findings link TDP-43 phase dynamics and aggregation to nuclear transport and other cellular processes.

## Results

### A chemical genetic screen identified modulators of TDP-43 phase separation

To identify cellular pathways modulating TDP-43 phase behavior, we conducted a chemical genetic screen using a small molecule library targeting diverse known cellular pathways and processes. We employed a previously established DLD1 cell model stably expressing an RNA-binding defective TDP-43 mutant (2KQ) tagged with the green fluorescence protein Clover, because this mutant undergoes rapid demixing upon induced expression (Yu et al., 2021), generating 20-30 sphere objects named anisosome in each nucleus. Both the number and size of anisosome increase substantially in the initial phase of TDP-43 induction but remain largely unchanged after 24 h of induction. We seeded cells in 384-well plates and induced the expression of TDP-43 2KQ with doxycycline for 24 hours. We then treated these cells with a LOPAC library containing 1280 drugs for 24 hours (Figure 1A). This approach could avoid false positive drugs that affect the expression of TDP-43 2KQ.

**Figure 1.**
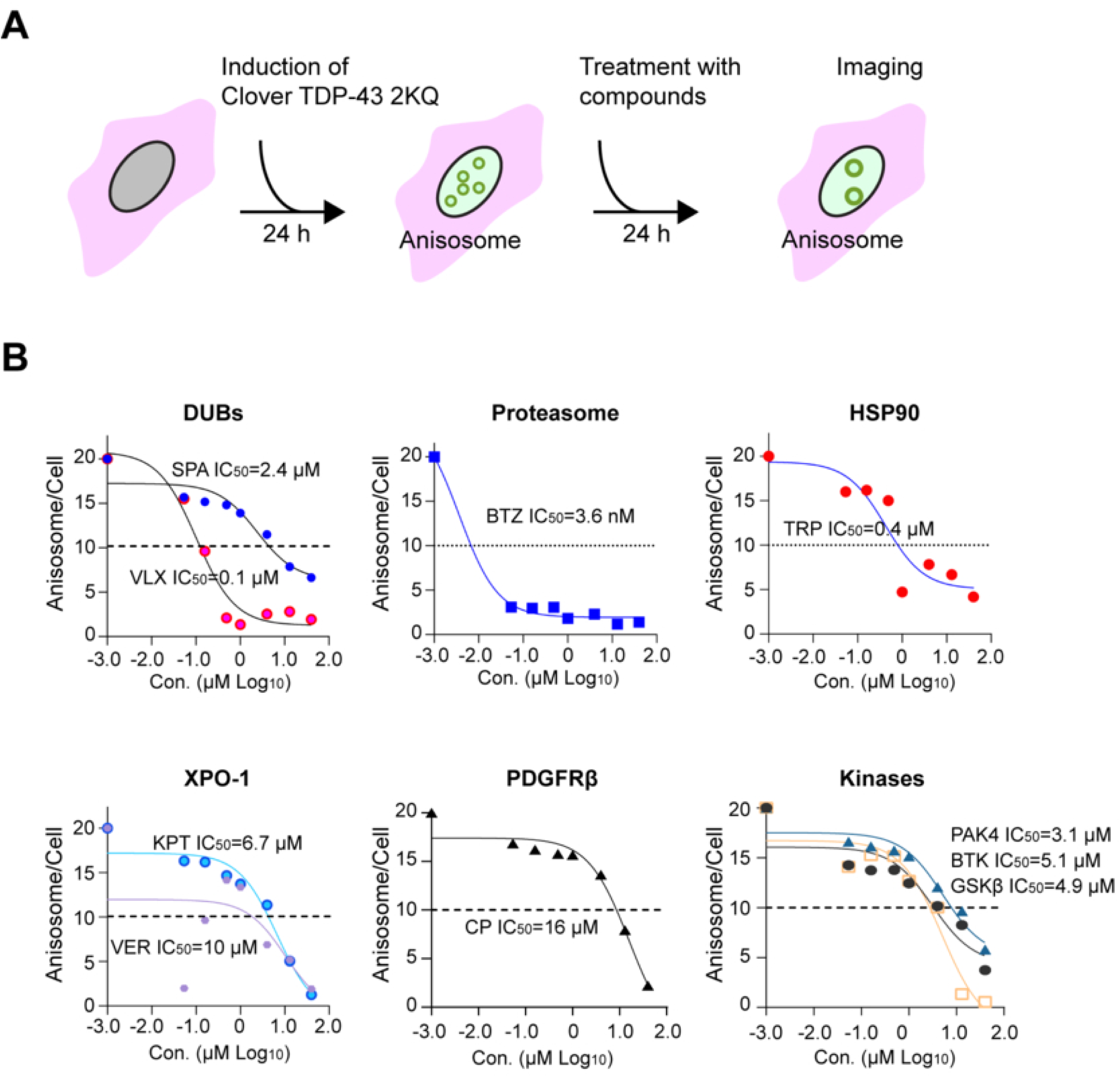
A chemical genetic screen identifies TDP-43 phase modulators. **(A)** Workflow of the chemical genetic screen. **(B)** Dose dependent reduction of anisosome number by identified chemicals. DLD1 TDP-43 2KQ-Clover cells were treated with doxycycline to induce anisosome for 24 h and then treated with drugs at the indicated concentrations. At least 500 cells were imaged for anisosome count per condition. A non-linear regression model was used to fit the data by GraphPad Prism.

High content imaging combined with algorithm-based image analyses (see methods) detected ∼20 spherical anisosomes per cell in controls and most treated cells. However, in cells treated with a subset of the drugs, the number of anisosomes was noticeably reduced (Supplemental Table 1). Importantly, cells treated with Bortezomib, a potent proteasome inhibitor, had fewer but enlarged TDP-43 positive puncta with irregular shapes (Figure 1B) similarly to cells treated with another proteasome inhibitor (van Eersel et al., 2011; Yu et al., 2021). This result validated our screening design.

To further confirm the identified hits, we performed a concentration titration experiment, which showed that these inhibitors dose-dependently reduced the anisosome number with IC_50_ ranging from sub-micromolar to micromolar (Figure 1B, Supplemental Table 1). Immunoblotting showed that treatment with these drugs did not change the total TDP-43 protein level (Supplementary Figure 1), suggesting that the effect on anisosome is not due to altered protein expression. Interestingly, the screen identified cytoplasmic signaling kinases including PAK4, BTK, and GSK-3β as well as the receptor tyrosine kinase PDGFRβ as modulators of TDP-43 phase behavior (Figure 1B, Supplemental Table 1). PAK4 and GSK-3β are known regulators of the YAP signaling pathway (Jin et al., 2025), and YAP was recently identified as a TDP-43-interacting protein that influences its phase dynamics (Zhang et al., 2025a). These connections provide additional physiological validation for the uncovered modulators. Our screen also links TDP-43 2KQ demixing to protein translation regulation because cycloheximide, a ribosome elongation inhibitor also affects TDP-43 phase dynamics (see below).

### Live cell imaging revealed two distinct classes of TDP-43 phase modulators

We next used confocal microscopy to further characterize selected inhibitors. We chose Tripterin (a drug targeting the heat shock protein HSP90), CP-673451 (a PDGFRβ inhibitor), two deubiquitinase inhibitors VLX-1570 and Spautin-1, and the Exportin-1 inhibitor KPT-276 because previous reports have linked the targets of these compounds to TDP-43 proteinopathies (Archbold et al., 2018; Farrawell et al., 2020; Furukawa et al., 2015; Garcia-Toscano et al., 2024). We first induced anisosome formation for 24 hours and then treated cells with these inhibitors. Compared to DMSO-treated controls, cells treated with Spautin-1 had fewer and smaller anisosomes. KPT-276 treatment also reduced anisosome number, albeit to a lesser extent (Figure 2A, top panels). In contrast, cells treated with CP-673451, Tripterin, or Bortezomib had fewer but larger TDP-43 puncta. Since Tripterin was also reported as a 20S proteasome inhibitor, we confirmed the involvement of HSP90 in this process using the well-established HSP90 inhibitor Geldanamycin (Supplemental Figure 2A). These results suggest that TDP-43 phase dynamics is modulated by the cellular folding capacity, the ubiquitin proteasome system, and nucleocytoplasmic transport.

**Figure 2.**
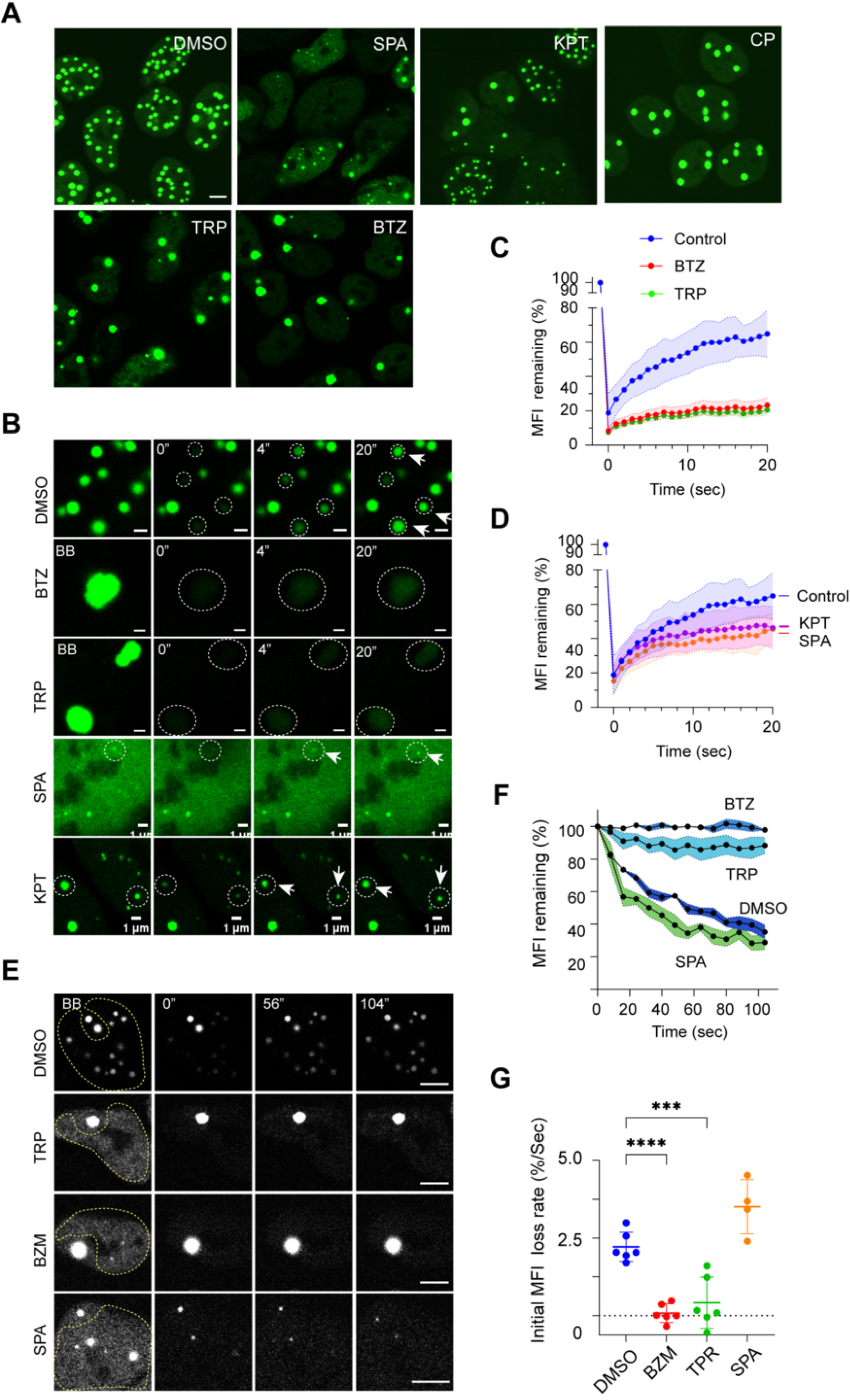
Two distinct types of TDP-43 phase modulators. **(A)** DLD1 TDP-43 2KQ-Clover cells treated with doxycycline for 24 h were further treated with the indicated compounds for 6 h and imaged (SPA, Spautin-1, 5 μM; KPT, KPT-276, 15 μM; CP, CP-673451, 30 μM; TRP, Tripterin, 1 μM; BTZ, Bortezomib, 10 nM). Shown are single confocal z-section views. Scale bar, 5 μm. **(B)** FRAP analyses of anisosomes after treatment with the indicated drugs for 3-5 h. Circles indicate bleached areas. Shown are single confocal z-section views. Scale bars, 1 μm. **(C, D)** Quantification of the experiments represented in B. MFI, mean fluorescence intensity. N = 5 anisosomes/condition. **(E)** Reverse FRAP analyses of anisosome dynamics. The areas indicated by the dashed lines were bleached. Shown are single confocal z-section views. Scale bar, 5 μm. **(F)** Quantification of the mean fluorescence intensity (MFI) loss over time in unbleached anisosomes as shown in E. 4-6 anisosomes were analyzed for each condition. **(G)** Quantification of the initial rate of fluorescence loss of unbleached anisosomes as shown in E. ****, p<0.0001; ***, p<0.001 by unpaired Student’s t-test (two-tailed). The rate of fluorescence loss in the first 10 seconds after photobleaching was calculated. 4-6 anisosomes were analyzed for each condition.

To further dissect the effects of these inhibitors on TDP-43 phase separation, we combined drug treatment with live cell fluorescence imaging, using Fluorescence Recovery After Photobleaching (FRAP) or reversed FRAP. In FRAP, when the green fluorescence of anisosomes was bleached by a laser in untreated cells, it rapidly recovered due to the recruitment of TDP-43 2KQ from the nucleoplasm (Figure 2B, top panels, Figure 2C, D). Conversely, in reversed FRAP, bleaching TDP-43 2KQ surrounding an anisosome repeatedly over time caused a gradual decline in fluorescence intensity within the unbleached anisosome, while the surrounding areas regained fluorescence partially (Figure 2E, top panels). This result suggests that TDP-43 2KQ in anisosomes undergoes rapid exchange with a pool in the nucleoplasm. Likewise, in cells treated with Spautin-1 or KPT-276, anisosome-associated TDP-43 2KQ could freely exchange with the nucleoplasmic pool (Figure 2B, D). In contrast, in cells treated with Bortezomib, Tripterin or Geldanamycin, the enlarged TDP-43 puncta appeared static; TDP-43 2KQ fluorescence within anisosomes failed to recover after photobleaching (Figure 2B, C, Supplemental Figure 2B), nor did it decrease over time when surrounding TDP-43 2KQ was bleached (Figure 2E-G). Collectively, these results suggest that these inhibitors modulate anisosomes via two distinct mechanisms: some maintain TDP-43 in a demixed liquid state, while others convert anisosomes to a gel-like solid state.

### A genome-wide siRNA screen identified genetic modulators of TDP-43 phase separation

To further decipher the molecular determinants of TDP-43 phase dynamics, we conducted an unbiased genome-wide siRNA knockdown (KD) screen (Figure 3A). To this end, we transfected a siRNA library targeting 21,404 human genes each with three siRNAs into DLD1 cells stably expressing Clover-tagged TDP-43 2KQ. After gene KD, anisosomes were induced for 24 hours. High content imaging and automated analysis identified 1,533 candidates whose KD reduced the anisosome number per cell (Z-score > 2) (Supplemental Table S2). To further narrow down the list, we performed a STRING protein network analysis based on the assumption that a protein interaction network bearing multiple positive hits would be more likely a true effector. We selected 211 networked genes with top Z-scores and re-screened each of them with three additional siRNAs. The result suggested the involvement of 110 genes in TDP-43 anisosome regulation (Figure 3A, Supplementary Table S3). GO pathway analyses categorized these genes into several pathways including RNA splicing, protein translation, proteasomal degradation, and nuclear transport (Figure 3B). The identification of ribosomal proteins and proteasome subunits is consistent with our chemical genetic screen, which has implicated translation and proteasomal degradation in anisosome modulation (Supplemental Table 1). Interestingly, GO analyses using ‘molecular function’ linked many identified genes to neurodegenerative diseases, particularly ALS (Figure 3C, D).

**Figure 3.**
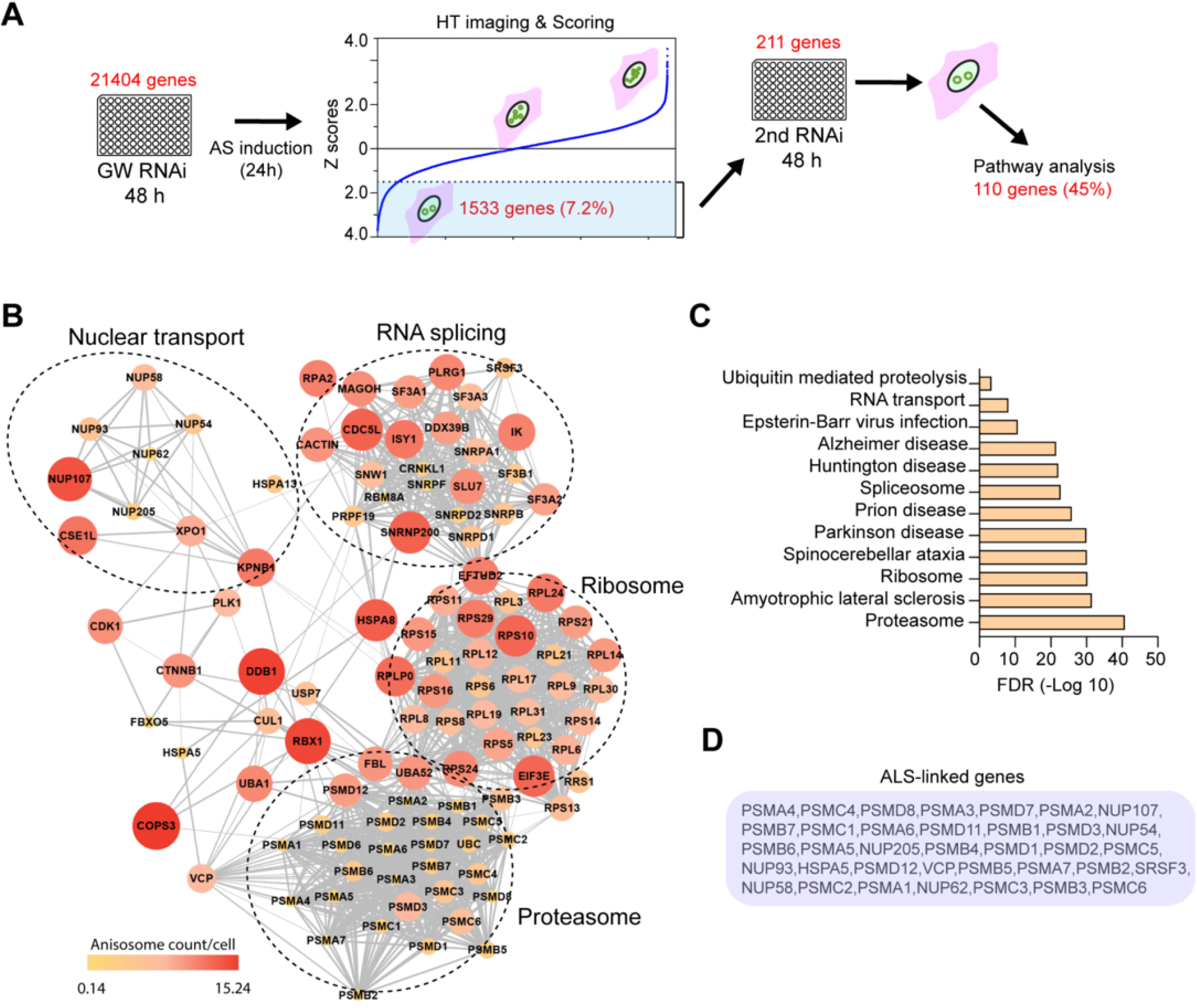
A genome-wide siRNA screen identifies modifiers of TDP43 phase behavior. **(A)** The workflow of the siRNA genetic screen. GW, genome-wide; AS, anisosome; HT, high-throughput. **(B)** Pathway analysis of genes whose knockdown reduces anisosome number in cells. The relative anisosome count are indicated by both color and size of the nodes. Candidate genes were selected based on Z-score >2 after normalization to the siRNA negative control. **(C)** GO molecular function analysis of TDP-43 phase modifiers. **(D)** A list of identified genes linked to ALS in C.

### Anisosome dynamics is modulated by RNA splicing and protein translation

Given the well-established function of TDP-43 in RNA binding and processing, we investigated the role of RNA splicing in anisosome regulation. To this end, we first incubated TDP-43 2KQ cells with doxycycline to induce anisosomes and then treated cells with a potent RNA splicing inhibitor Pladienolide-B (PlaB). Confocal microscopy confirmed that RNA splicing inhibition resulted in fewer but larger TDP-43-positive puncta in a dose dependent manner (Figure 4A, B, Supplemental Figure 3A).

**Figure 4.**
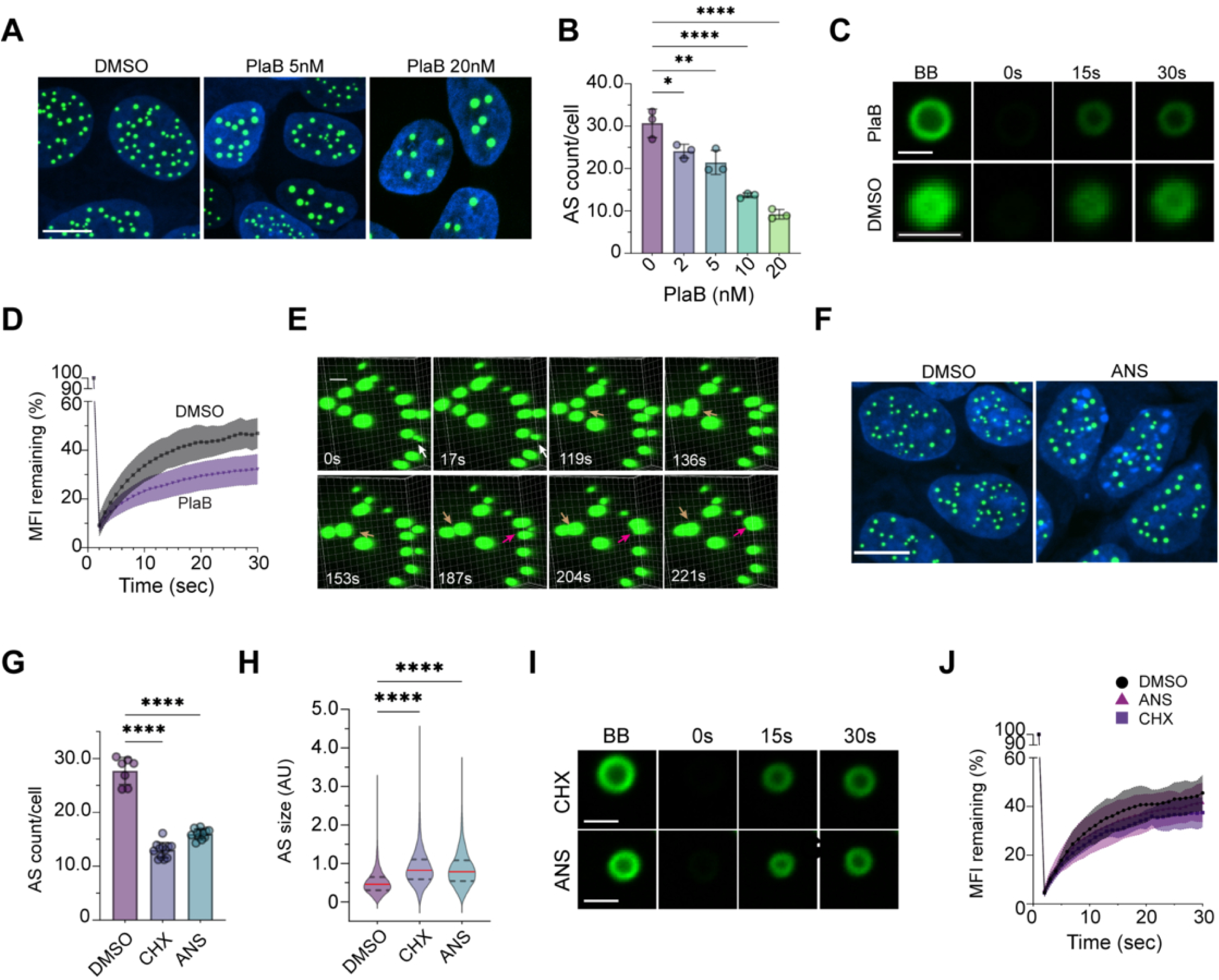
Anisosome phase behavior is modulated by RNA splicing and protein translation. **(A)** The splicing inhibitor Pladienolide-B (PlaB) reduces anisosome number in a dose dependent manner. Representative images of anisosome-induced cells treated with DMSO (control), 5 nM, or 20 nM PlaB for 16 h. Shown are maximum intensity projection views. Scale bar: 10 µm. **(B)** Quantification showing the number of anisosome (AS) per cell in TDP-43 2KQ-Clover cells treated with PlaB as indicated. * p <0.05, ** p <0.01, **** p < 0.0001 by ordinary one-way ANOVA and Dunnett’s multiple comparisons. n=3 independent experiments, and in each experiment, at least 30 randomly selected cells were analyzed. **(C)** Representative z-section confocal FRAP images of anisosomes in cells treated for 16 h with DMSO or PlaB (20 nM). BB, before photobleaching, right after photobleaching (0 s), or 4 and 30 seconds after photobleaching (4 s and 30 s). Scale bar, 1 µm. **(D)** The graph shows the quantification of the remaining TDP-43 fluorescence (FL) in C. Error bars indicate mean ± SD, N = 28 for control and 23 for PlaB-treated cells. MFI, mean fluorescence instensity. **(E)** Live cell imaging of anisosome fusion in TDP-43 2KQ-Clover cells treated with 20 nM PlaB for 5 h before tracking the fusion. Representative reconstructed 3-D images from Movie S1 showing fusion events indicated by arrows. Scale bar, 1 µm. **(F)** Representative maximum intensity projection views of anisosome-induced (24 h) cells treated with DMSO (control) or ANS (200 nM) for 16 h. Scale bar, 1 µm. **(G)** Quantification of the number of anisosomes per cell in randomly selected images of DMSO-, Cycloheximide (CHX)-, or ANS-treated cells. **** p < 0.0001 by ordinary one-way ANOVA and Dunnett’s multiple comparisons. Each dot represents a randomly selected field with at least 20 cells counted from one of the 3 independent experiments. **(H)** Quantification of anisosome size in control or cells treated with CHX or ANS. ****, p < 0.0001 by ordinary one-way ANOVA and Dunnett’s multiple comparisons. N=3 independent experiments. AU, arbitrary unit. **(I)** As in **C** except that anisosome-induced cells were treated with CHX or ANS before photobleaching. Scale bar 1 µm. **(J)** Quantification of fluorescence recovery in CHX- or ANS-treated cells vs the DMSO control. N=10 anisosomes/condition.

Despite size difference, anisosomes in drug-treated cells were morphologically indistinguishable from controls. FRAP experiments further showed that TDP-43 within PlaB-treated condensates remained highly mobile, although the fluorescence recovery rate was slightly reduced compared to controls (Figure 4C, D). Time-lapse live cell imaging frequently detected fusion of TDP-43 puncta after PlaB treatment (Figure 4E, Movie S1). Together, these results suggest that disrupting RNA splicing stabilizes TDP-43 in a demixed liquid state, resulting in larger condensates.

Next, we explored the effect of translation inhibition on anisosome dynamics using Cycloheximide (CHX) and Anisomycin (ANS), two translation elongation inhibitors. Confocal imaging and immunoblotting showed that translation inhibition by CHX or ANS did not significantly reduce TDP-43 expression (Supplemental Figure 1, 3). Like PlaB-treated cells, CHX- and ANS-treated cells contained fewer, enlarged anisosomes (Figure 4F-H, Supplemental Figure S2B). FRAP analysis revealed that TDP-43 dynamics within anisosomes were similar between ANS-treated and control cells (Figure 4I, J). These results suggest that translation inhibition also stabilizes TDP-43 in liquid condensates.

### Nuclear export modulates TDP-43 liquid-to-solid phase transition

Our screen also identified several nuclear transport regulators (e.g., Exportin-1/XPO1, Exportin-2/CSE1L) and nuclear pore components as anisosome modulators (Figure 3B). We focused on XPO1 because our chemical genetic screen identified two XPO1 inhibitors, KPT-276 and Verdinexor as potent anisosome modulators (Figure 1B).

To explore the role of XPO1 in TDP-43 anisosome regulation, we induced anisosome formation in Clover-TDP-43 2KQ cells and treated them with Leptomycin B (LMB), a potent XPO1 inhibitor. LMB treatment resulted in a time- and dose-dependent reduction in anisosome number, while increasing their sizes (Figure 5A-C). The enlarged TDP-43 puncta showed the typical hollowed anisosome ring structure (Figure 5D), indicating that TDP-43 remained in a demixed liquid state. Indeed, FRAP experiments demonstrated that LMB treatment did not significantly affect the fluorescence recovery rate of TDP-43 within bleached anisosomes (Figure 5D, E). Time-lapse confocal imaging showed that preformed anisosomes began to fuse with each other 5 hours after LMB treatment (Figure 5F, Movie S2). Thus, XPO1 inhibition stabilizes TDP-43 in a liquid phase.

**Figure 5.**
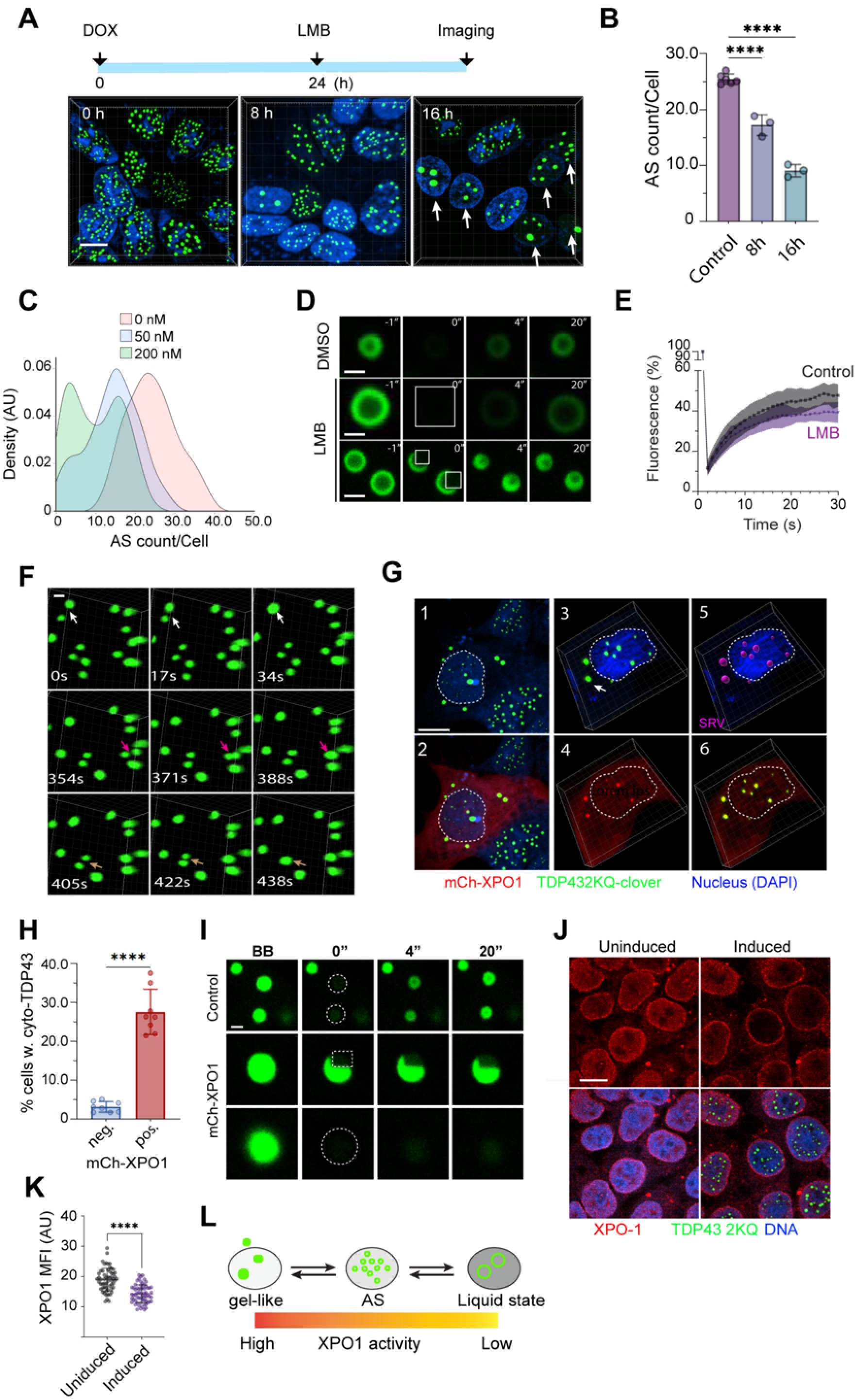
XPO1 regulates anisosome liquid-to-solid transition. **(A)** Pharmacological inhibition of XPO-1 with Leptomycin B (LMB, 200 nM) reduces the number of anisosome. The graph on top indicates the experimental design. Arrows show cells with enlarged anisosomes. Scale bar 10 µm **(B)** Quantification of anisosome numbers per cell in experiments represented by A. Error bars indicate mean ± SD; **** p <0.0001 by ordinary one-way ANOVA and Dunnett’s multiple comparisons. n=3 independent experiments, each with duplicated controls. **(C)** LMB dose dependently reduces anisosome number. Anisosome were induced in TDP-43 2KQ-Clover cells followed by treatment with LMB at the indicated concentrations for 16 h. The histogram shows the distribution of cells (50-90 cells/condition) by anisosome count. **(D)** FRAP experiments demonstrate that anisosomes remain in a liquid phase following LMB treatment (200 nM, 16 h). Anisosome-induced (24 h) cells were treated with DMSO or LMB for 16 h and then photobleached at the indicated areas. Scale bar 1 µm. **(E)** Quantification of the FRAP experiment in D. N=18 for control and 16 anisosomes for LMB-treated cells. **(F)** Time-lapse confocal microscopy detects anisosome fusion after LMB (200 nM, 5 h) treatment in TDP-43 2KQ-Clover cells. Arrows indicate fusion events. Scale bar, 1 µm **(G)** Overexpression of mCherry-tagged XPO-1 in TDP-43 2KQ-Clover cells induces cytoplasmic TDP-43 puncta. TDP-43 2KQ-Clover cells (green) transfected with mCherry-XPO1 (red) were stained with DAPI (blue) to label nuclei (dashed lines). Cells were imaged 48 h post-transfection. Panels 1, 2 show a representative confocal section, while panels 3-6 show reconstructed 3-D views. The position and volume of anisosomes were also presented in magenta in a surface-rendered view (SRV) in panel 5. The arrow in panel 3 indicates an example of cytoplasmic TDP-43 aggregate. Scale bar, 10 µm. **(H)** Quantification of the percentage of cells showing cytosolic TDP-43 puncta in XPO-1 positive (pos) and negative (neg) cells in 8 randomly selected fields each with at least 20 cells from 3 independent experiments. **** p <0.0001 by two-tailed unpaired Student’s t-test. **(I)** FRAP-based confocal imaging reveals the transition of anisosomes into a gel-like state upon XPO-1 overexpression. Scale bar, 1 µm. **(J)** Anisosome formation changes endogenous XPO-1 localization. TDP-43 2KQ-Clover before or after anisosome induction were stained with anti-XPO-1 antibodies (red) and DAPI (blue). The bleached areas were indicated by dashed lines. Scale bar, 10 µm. **(K)** Quantification of nuclear endogenous XPO-1 mean fluorescence intensity (MFI) in individual cells (indicated by dots) before or after anisosome induction. **** p < 0.001 by two-tailed unpaired Student’s t-test. N = 58 cells for uninduced and 63 cells for induced condition in 2 independent repeats. **(L)** A schematic model depicting how the nuclear XPO-1 activity influences TDP-43 liquid-to-phase transition. AS, anisosome.

To corroborate the inhibitor study, we overexpressed mCherry-tagged XPO1, a condition known to enhance nuclear export of XPO1 cargos (Antonarakis, 2018; Kim et al., 2022; Tang et al., 2025). Interestingly, we observed enlarged TDP-43 positive puncta in mCherry-XPO1-expressing cells with characteristics distinct from that of XPO1-inhibited cells (Figure 5G, panels 1, 2). First, we did not observe the typical hollowed anisosome ring structures in mCherry-XPO1-expressing cells. Instead, these large puncta appeared to have irregular shapes. Secondly, while anisosomes in mCherry-XPO1 negative cells were almost exclusively nuclear, ∼30% of mCherry-XPO1-expressing cells contained cytoplasm-localized TDP-43 puncta (Figure 5G, panels 1, 2, 3, Figure 5H). Thirdly, FRAP experiments demonstrated that Clover-TDP-43 fluorescence in mCherry-XPO1-expressing cells did not recover after photobleaching, suggesting that TDP-43 was in a gel-like state in transition to aggregates (Figure 5I). As expected, over-expression of mCherry did not alter the subcellular localization of TDP-43 puncta (Supplemental Figure S4). Thus, higher XPO1 activity is associated with increased accumulation of TDP-43 in a gel-like form in the cytoplasm.

Immunostaining of XPO1 showed that anisosome induction depleted endogenous XPO1 from the nucleoplasm (Figure 5J, K). In contrast, no XPO-1 accumulated within anisosomes, probably because antibodies could not stain proteins inside anisosomes due to poor accessibility (Yu et al., 2021).

Consistent with this interpretation, we noticed the presence of over-expressed mCherry-XPO1 in TDP-43 puncta in some cells (Figure 5G, panels 3-6). Since it is known that XPO1 cannot bind TDP-43 directly (Pinarbasi et al., 2018), our data suggested a possible indirect interplay between TDP-43 and XPO1.

However, the precise mechanism requires future elucidation. Collectively, our results suggest that the stability and dynamics of anisosomes are modulated by XPO1; Low XPO1 activity stabilizes TDP-43 in a liquid state and favors larger nuclear condensates, while high XPO1 activity is associated with a transition to gel-like structures (Figure 5L).

### Nuclear export inhibition stabilizes TDP-43 anisosomes in an RNA dependent manner

To dissect the factor(s) mediating the effect of nuclear export on TDP-43 phase regulation, we developed a semi-permeabilized cell–based *in vitro* assay (Figure 6A). To this end, anisosomes were first induced in Clover-tagged TDP-43 2KQ cells, followed by selective permeabilization of the plasma membrane using the pore-forming toxin streptolysin O (SLO). Permeabilization was monitored at 37 °C by confocal microscopy using the membrane-impermeable dye NucSpot, which stained nuclei only upon plasma membrane rupture. Notably, cell permeabilization led to a marked reduction in TDP-43–positive nuclear puncta and the overall fluorescence intensity (Figure 6B). Quantitative analysis revealed an inverse correlation between the NucSpot signal and TDP-43 fluorescence intensity (Figure 6C).

**Figure 6.**
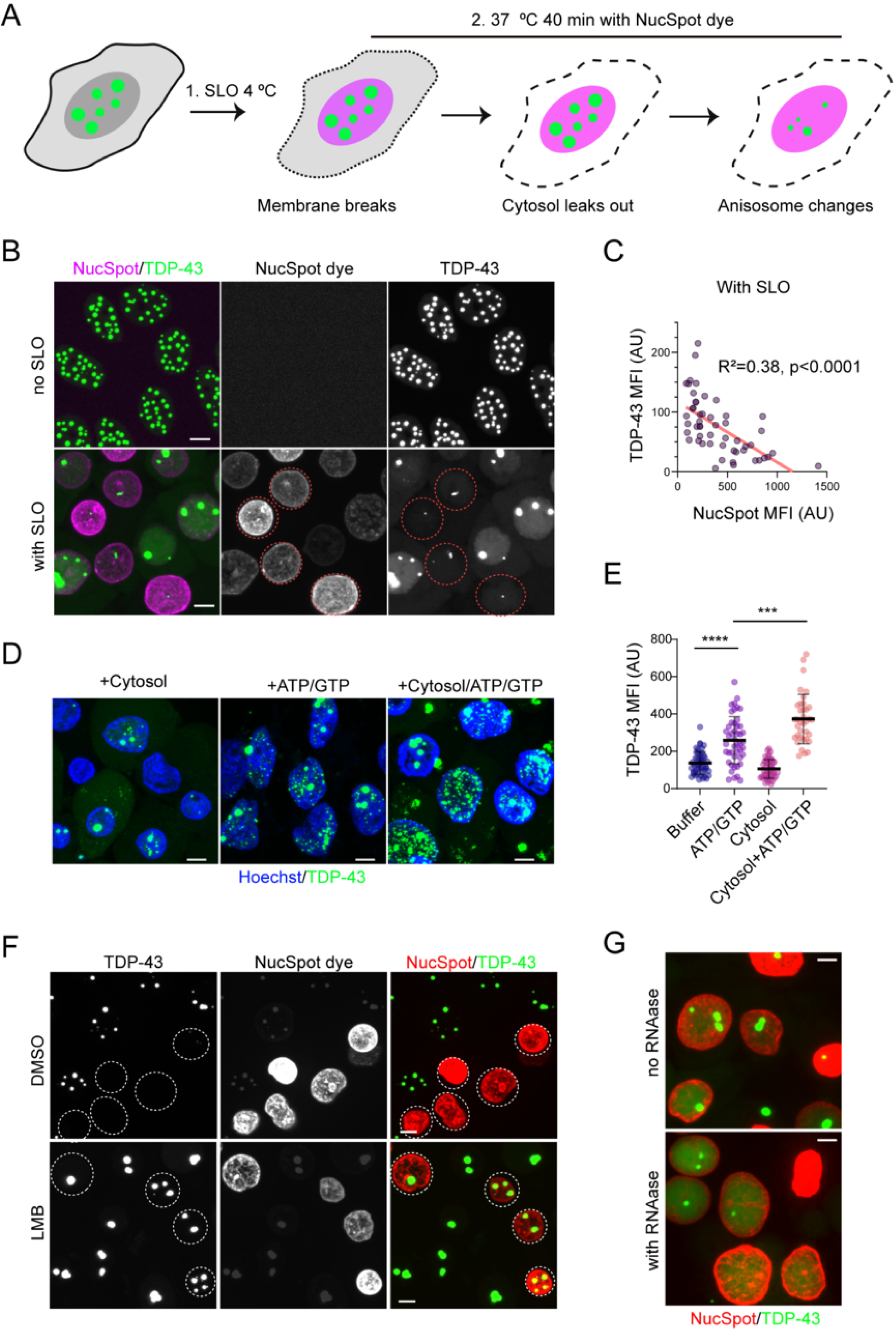
The inhibition of XPO-1 stabilizes anisosomes in an RNA-dependent manner. **(A)** A schematic diagram of the semi-permeabilized cell assay. SLO, streptolysin O. **(B)** TDP-43 in anisosomes becomes soluble after cell permeabilization (indicated by NucSpot positive staining in magenta). Note: The intensity of NucSpot dye correlates with the permeabilization time. Dashed circles highlight cells permeabilized early during the incubation. The cells have fewer anisosomes. Scale bars, 5 μm. **(C)** TDP-43 mean fluorescence intensity (MFI) displays an inverse correlation with the NucSpot dye intensity in B. AU, arbitrary unit. N =52 cells. **(D)** Small TDP-43 positive puncta were reformed after the incubation of permeabilized DLD1 cells with cow liver cytosol in the presence of ARS and GTP (100 μM) (ATP/GTP) at 37 ℃for 40 min. Cells incubated with cytosol without ATP/GTP or with PBS plus ATP/GTP serve as controls. Cells were fixed and stained with Hoechst (blue) to label nuclei. Scale bars, 5 μm. **(E)** Quantification of the TDP-43 mean fluorescence intensity (MFI) in cells as shown in D. Error bars, SD. ****, p<0.0001; ***, p<0.001 by unpaired Student’s t-test. N=at least 35 randomly selected cells representing two independent experiments. **(F)** LMB treatment (200 nM, 16 h) stabilizes anisosomes. Dashed circles indicate permeabilized cells. Scale bars, 5 μm. **(G)** RNAase T1 treatment destabilizes anisosomes in DKD1 cells pre-treated with LMB (200 nM, 16 h). DLD1 cells induced for TDP-43 expression were treated with LMB (200 nM, 16 h). Cells were permeabilized in the absence or presence of RNAase T1 for 40 min before imaging. Images shown in this figure are maximum intensity projection views of confocal z-sections. Scale bars, 5 μm.

The reduction in TDP-43 signal probably resulted from a shift of TDP-43 from a phase-separated high fluorescent state into a soluble state with reduced fluorescence intensity (Zhang et al., 2026). We attributed this phenotype to the depletion of cytosolic factors and ATP during cell permeabilization because it is known that anisosome formation and maintenance require HSP70, a cytosolic ATPase (Yu et al., 2021). To test this idea, we incubated permeabilized cells with purified cytosol together with an ATP-regenerating system (ARS) and GTP to reconstitute a condition resembling intact cells. Indeed, we observed reformation of TDP-43 puncta after incubation at 37 ℃ (Figure 6D). Although the puncta were smaller than anisosomes, their formation significantly increased TDP-43 fluorescence intensity (Figure 6E). Addition of buffer with ARS and GTP modestly increased TDP-43 puncta and fluorescence intensity, while cytosol alone had no effect (Figure 6D, E). Although the size limitation prevented further characterization of these puncta, the fact that their formation depends on energy and cytosolic factors is consistent with the requirement of HSP70 in anisosome formation (Yu et al., 2021).

We next applied this assay to examine the mechanism underlying LMB-induced enlargement of anisosomes. Strikingly, anisosomes in LMB-treated cells remained stable following permeabilization (Figure 6F). However, inclusion of RNase T1, which degrades single-stranded RNA during permeabilization led to their dissolution (Figure 6G). These findings suggest that LMB stabilizes anisosomes at least in part by increasing nuclear RNA availability.

### Nuclear export inhibition mitigates TDP-43 hyperphosphorylation in an organoid model of TDP-43 proteinopathy

Since XPO1 inhibition causes TDP-43 to accumulate in a demixed liquid state, we asked whether reducing XPO1 activity could mitigate TDP-43 cytosolic phosphorylation. To test this idea, we used a recently established organoid model of TDP-43 proteinopathies (Zhang et al., 2025b), which is derived from an induced pluripotent stem cell (iPSC) line carrying the ALS-associated mutation (K181E) in the endogenous TDP-43 locus (Zhang et al., 2025b). Like the 2KQ mutant, the K181E substitution disrupts the RNA binding activity of TDP-43 (Chen et al., 2019). Additionally, the K181E mutant was reported to accumulate in a condensed liquid phase in association with HSP70 in cultured cells under overexpression conditions, and like anisosomes, these condensates can be converted to solid aggregates during cellular stress or HSP70 depletion (Gu et al., 2021).

We treated organoids bearing the endogenous K181E mutation or wild-type (WT) controls with KPT-276 at a low dose (20 nM) for 35 days starting on day 87 based on pilot data indicating good tolerance with prolonged treatment. Homozygous mutant organoids were used to facilitate phosphor-TDP-43 detection. We sectioned and stained these organoids with antibodies against Serine 409/410-phosphorylated TDP-43 (p-TDP-43) and total TDP-43 since hyperphosphorylated TDP-43 (S409/410) is a clinically validated pathological marker for TDP-43-containing cytoplasmic aggregates. Confocal imaging detected significant p-TDP-43 signal in mutant but not in WT organoids (Figure 7A, B), as reported previously (Zhang et al., 2025b). By contrast, we did not detect an increase in total TDP-43 staining (green) in mutant organoids (Figure 7C). Interestingly, KPT-276 treatment significantly decreased p-TDP-43 signal in K181E organoids without affecting the total TDP-43 signal (Figure 7A, B, C). As expected, mutant organoids had more cells with reduced nuclear TDP-43 signal compared to WT organoids (Figure 7A, right panels, D). KPT-276 treatment significantly rescued this phenotype (Figure 7D). These results imply that maintaining TDP-43 in a nuclear demixed liquid state mitigates p-TDP-43 accumulation but whether the reduction of pTDP-43 immunoreactivity reflects a decrease in insoluble TDP-43 aggregates remains to be determined.

**Figure 7.**
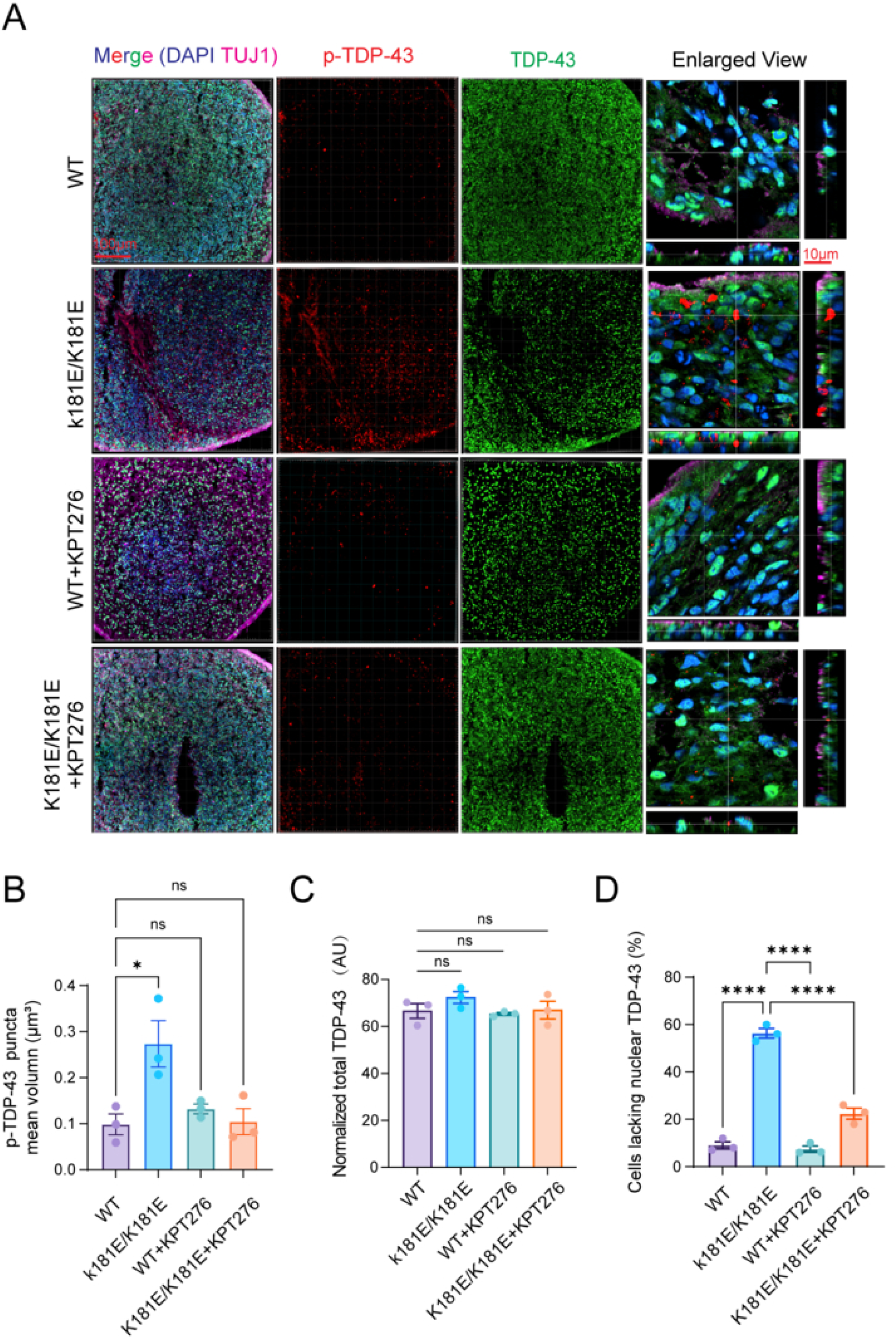
Inhibition of XPO-1 mitigates TDP-43 hyperphosphorylation in TDP-43 K181E organoids. **(A)** Confocal fluorescence imaging reveals significant reduction in phosphorylated TDP-43 (p-TDP43) in K181E/K181E organoids following KPT-276 treatment. Organoids of the indicated genotypes at day 87 were treated with DMSO as a control, or with KPT-276 (20 nM) for 35 days. Organoids were fixed, sectioned, and stained with antibodies against TUJ1, a neuronal marker (Magenta), TDP-43 (green) and p-TDP-43 (red). n= 3-5 organoids from two individual batches. **(B, C, D)** Quantification of p-TDP-43 puncta mean volume (B), total TDP-43 intensity (C) and the percentage of cells with reduced nuclear TDP-43 (D) from experiments represented by **A**. For B, p-TDP-43-positive puncta were segmented from collected 3-D images for mean puncta volume calculation. Error bars indicate mean ± SEM. * p < 0.05, **** p < 0.0001, ns, not significant, by one-way ANOVA. n = 3 organoids per condition.

## Discussion

ALS- and FTD-associated TDP-43 mutants defective in RNA binding are known to undergo self-association through their intrinsically disordered regions and RNA-binding domains, generating demixed liquid, gel, and aggregated states in dynamic exchange. This protein phase behavior is shared by many RNA binding proteins and is increasingly recognized as a contributor to protein aggregation in neurodegenerative diseases (Shorter, 2019). Intriguingly, TDP-43 phase separation can form unique layered nuclear structures named anisosome. Although it is unclear how these structures are nucleated in the cell, our study suggests that, once formed, anisosomes can grow by recruiting additional TDP-43 from the surrounding environment. Small anisosomes may also fuse with each other, although such fusion events become less frequent after anisosomes mature.

Our study supports the notion that TDP-43, at high concentrations, can change from a demixed liquid state into a gel-like state en route to amorphous aggregation (Babinchak et al., 2019; Carey and Guo, 2022). We found that proteasome and HSP90 inhibition or enhancing XPO-1-mediated nuclear egression promotes the conversion of TDP-43 anisosomes into a gel-like state. By contrast, inhibition of RNA splicing, protein translation, or nuclear export all favors a liquid state.

Our validation of proteasome inhibition as a driver of TDP-43 droplet enlargement and hardening reinforces the idea that impaired protein clearance promotes the transition of TDP-43 from liquid condensates to gel-like or solid aggregates. This is consistent with previous studies in ALS models, which showed that proteasome stress accelerates TDP-43 aggregation and toxicity (van Eersel et al., 2011). In contrast, inhibition of deubiquitinases reduced the number of TDP-43 anisosomes, probably by promoting the proteasome-mediated clearance of TDP-43. The role of HSP90 in this process may be linked to TDP-43 folding or stability, as HSP90 can bind TDP-43 to prevent promiscuous interactions (Lin et al., 2021). Collectively, these observations suggest that TDP-43 condensates are maintained by a delicate balance between formation and clearance, modulated by proteolytic and chaperone activities.

While our chemical and genetic screens have identified multiple pathways influencing TDP-43 phase states, a particularly intriguing aspect of our study is the discovery that nuclear transport modulates the TDP-43 liquid-to-solid transition. However, given the lack of detectable interaction between XPO-1 and TDP-43 as suggested previously (Duan et al., 2022; Pinarbasi et al., 2018), XPO1 probably modulates TDP-43 phase behavior via an indirect mechanism. Indeed, our experiments using semi-permeabilized cells suggest that nuclear transport might affect TDP-43 phase behavior by altering the abundance of nuclear RNA. Anisosome maturation might involve one or more mRNA splicing intermediates. If so, the completion of mRNA splicing followed by nuclear egression of spliced RNAs would disfavor anisosome growth and maturation. However, our data does not exclude other RNA species or RNA-binding proteins as anisosome stabilizer. Whether RNA-dependent stabilization of anisosomes operates in the same way in intact cells also requires further validation. Nevertheless, our findings hint at a potential feedback mechanism in which TDP-43 demixing perturbs nuclear export, analogous to how cytoplasmic TDP-43 aggregates disrupt nuclear import.

Because the TDP-43 2KQ mutant lacks RNA binding activity, anisosome maturation is likely modulated by other RNA-binding factors. In this regard, TDP-43 is known to interact with many splicing factors including members of the hnRNP family like hnRNP A1/A2 and PSF (Freibaum et al., 2010).

RNA splicing inhibition might alter TDP-43’s interactions with these factors, trapping it in an RNA-containing complex that favors anisosome growth and maturation, This model could explain why both PlaB and LMB reduce the number of TDP-43 anisosomes while maintaining their dynamic liquid property.

Together, our results support a model in which multiple cellular pathways such as proteostasis network, RNA metabolism, protein translation, and nuclear export cooperatively determine the phase behavior of TDP-43. These findings have deepened our understanding of how physiological phase separation can evolve into pathogenic aggregation through cumulative failures of cellular quality control and nuclear transport systems.

## Methods

### RNAi-based genetic screen for anisosome regulators

Genome-wide RNA interference (RNAi) screen was performed using DLD1 cells stably expressing Clover-tagged TDP-43 2KQ. Cells were transfected with siRNAs targeting 21,404 genes for 72 hours, followed by induction of anisosome formation with doxycycline (0.5 µg/mL) for 24 h. Specifically, the screen was carried out in 384-well format with 3 individual siRNAs for each gene. To prepare siRNA transfection, control siRNA (40 nmol) was mixed with 2 mL water to create a 50X (20 µM) stock and siRNA-death (20 nmol) was mixed with 1 mL of water for the same concentration. Dilute siRNA stocks to a 400 nM working concentration in water. For each well, mix 0.1 µL transfection reagent in 20 µL of cell culture medium without FBS or P/S. Add 2 µL of siRNA solution to each well and incubate with RNAiMAX for 30 minutes at room temperature. While incubating RNAiMAX, prepare trypsinized cells by ensuring thorough separation and accurate cell counting. Add 20 µL of cell suspension at a density of 0.43 × 10^5^ cells/mL (850 cells/well) in medium containing 20% FBS and 2 x P/S to achieve a final volume of 40 µL per well. Incubate cells for 3 days before addition of 10 µL 10 x doxycycline solution to each well (final volume 50 µL) for anisosome induction. 24 h later, high-throughput (HT) imaging and automated phenotypic screening were conducted to identify genes that modulated anisosome formation. To control for siRNA transfection efficiency, we routinely included a positive knockdown control in which cells were treated with a mixture of siRNAs targeting several essential genes (AllStars Hs Cell Death siRNA (Qiagen, #1027299).

Fluorescence imaging was conducted using an Opera Phenix High-Content Screening System (PerkinElmer). Whole-well images were acquired in confocal mode using a 20X water objective lens after anisosome induction for 24 h. Images were reconstructed and analyzed using the PerkinElmer Columbus server (v2.9.1). Nuclei were segmented based on Hoechst33342 staining, and anisosomes were detected via YFP signals within the nuclear region. The mean anisosome count per nucleus was calculated to assess changes in anisosome levels, while toxicity was evaluated by counting the total number of nuclei. Data were normalized to the siRNA-Neg control for each plate. To identify significant hits, Z-scores were calculated by subtracting each anisosome count from the plate sample mean. The resulting values were divided by the plate’s S.D. value. Identified 1,533 genes whose knockdown significantly reduced the number of anisosomes per cell (Z-score > 2) were subjected to protein network analysis by STRING to identify candidate anisosome regulatory genes that were subject to a second-round screen.

### A chemical genetic screen for anisosome regulators

We also performed a chemical genetic screen using LOPAC^R1280^ (Sigma) compound library on DLD1 cells expressing Clover-tagged TDP-43-2KQ. Briefly, cells were induced with doxycycline for 24 h to promote anisosome formation, followed by treatment with individual compounds for an additional 24 hours. High-throughput (HT) imaging and automated phenotypic screening were conducted to identify compounds that modulated anisosome formation.

### Visualization and analysis of anisosome dynamics

To examine the impact of drug treatment on anisosome size, cells were induced to form anisosome for 24 h and then treated by different inhibitors as follows: cycloheximide (CHX, 20 µg/mL), anisomycin (ANS, 200 nM) for 16 hours. For splicing inhibition, cells were exposed to Pladienolide-B (PlaB) at concentrations of 5 or 20 nM, or DMSO (control) for 16 hours before imaging. Cells were fixed with 4% paraformaldehyde in Phosphate Buffer Saline (PBS). Randomly selected fields were used to quantify the number of anisosomes per cell with at least two independent repeats.

To image anisosome fusion, live cell imaging was performed by treating cells with Leptomycin B (200 nM) or PlaB (20 nM) for 5 hours and then imaged by confocal 3-D sectioning for 30 minutes to capture anisosome fusion events. For Fluorescence Recovery After Photobleaching (FRAP), cells were treated with DMSO, Leptomycin B, Geldanamycin or PlaB as indicated in figure legends. Either part of an anisosome or entire anisosome was photobleached and imaged. TDP-43 fluorescence recovery after photobleaching was quantified by ImageJ and calculated to assess anisosome dynamics. At least 10 anisosomes for each condition were bleached and analyzed.

To evaluate the time course of Leptomycin B (LMB)-induced anisosome changes, DLD1 TDP-43 2KQ-Clover cells were seeded and induced with 0.5 µg/mL doxycycline for 24 hours. Cells were treated with 10 µM LMB and fixed at 0-, 8-, and 16-hours post-treatment. Fixed cells were stained with Hoechst to visualize nuclei. 10-20 confocal 3-D sections were obtained for each randomly selected field. Images were converted to maximum projected view by ImageJ before anisosome counting and intensity measurement. The density plot of anisosome counts per cell was generated using R, illustrating dose-dependent effects of leptomycin on anisosome formation.

To test the effect of XPO1 overexpression on TDP-43 phase regulation, DLD1 TDP-43 2KQ-Clover cells were co-transfected with mCherry-tagged XPO1 (mCh-XPO1) for 24 h before doxycycline was added at 0.5 µg/mL to induce anisosome formation for 24 h before confocal imaging.

To study the relationship between anisosome induction and endogenous XPO1 localization, cells were seeded, induced with 0.5 µM doxycycline for 48 hours, and fixed with paraformaldehyde in PBS. Immunostaining was performed using an XPO1 antibody (Cell Signaling, 46249S, 1:125) and Hoechst nuclear stain. Fluorescence intensity of XPO1 in both induced and uninduced cells was measured by ImageJ.

### Semi-permeabilized cell experiments

To reconstitute anisome disassembly and re-assembly in seminar permeabilized cells, 30,000 DLD1 cells were seeded in an eight-well ibidi cell chamber two days before the experiment. Cells were treated with Doxycycline for 24 h to induce anisosomes and then treated with inhibitors as indicated in the figure legends. We then washed cells three times with ice-cold PBS containing 2 mM MgCl_2_ (PBS-Mg) and then treated these cells in the same buffer (300 µl) containing 200 units of SLO (Sigma-Aldrich, cat. no. SAE0089) on ice for 30 min. The cells were then washed two times with PBS-Mg buffer and incubated with 200 µl of PBS buffer containing the NucSpot dye at 37 ℃ for 30-40min to permeabilize the cell and disassemble anisosomes. After that, cells were incubated with 200 µl cow liver cytosol (Lee et al., 2025; Ye et al., 2001) in the absence or presence of an ATP regenerating system (Ye et al., 2001) and GTP (100 μM) at 37 ℃ for 30-40 min. Cells were either imaged directly or fixed with 4% paraformaldehyde before imaging by a Nikon CSU-W1 SoRa spinning disk confocal microscope.

### Experiments with organoids

Stem cell gene editing of the patient K181E mutation in iPSC cells was reported previously (Zhang et al., 2025b). Differentiation of iPSCs into forebrain neurons was performed following a previously published protocol (Fernandopulle et al., 2018). Briefly, the procedure used an induction medium (IM-N2) and a neuronal culture medium (CM). IM-N2 was prepared with Knockout DMEM/F12, N2 supplement, NEAA, Gluta-MAX, Chroman I, and doxycycline. On Day 0, iPSCs were observed for confluency, dissociated with Accutase, centrifuged, and resuspended in IM-N2 with Chroman I, and then seeded into Matrigel-coated plates. Over the next four days, cells were monitored microscopically for neurite extensions while media containing doxycycline is refreshed daily. By Day 4, cells exhibit neurite growth and are ready for replating onto poly-L-ornithine (PLO)-coated dishes, prepared in advance by coating with PLO solution, incubating, washing, and drying. Replated cells were cultured in CM comprising BrainPhys medium, B27+ supplement, neurotrophic factors (GDNF, BDNF, NT-3), laminin, and doxycycline, to promote neuronal maturation.

3-D culture and organoid growth were performed as previously described (Nguyen, 2022). In brief, iPSCs grown on Matrigel were dissociated into single cell suspension by Versene solution (ThermoFisher) and seeded into a 12-well Aggrewell plate (Stemcell Technologies) at 4,000 cells/ well. Next day, spheroids were transferred to an ultralow attachment plate (Corning) containing phase I medium: DMEM/F12, 20% Gibco KnockOut Serum Replacement, 1X Glutamax, 1X MEM Nonessential Amino Acid, 55 µM β-Mercaptoethanol, 1X Pen/Strep, 2 µM Dorsomorphine, 2 µM A83-01. After 5 days, medium was switched to phase II medium: DMEM/F12, 1X N-2 Supplement, 1X Glutamax, 1X MEM Nonessential Amino Acid solution, 1X Pen/Strep, 4 ng/mL WNT3a, 1 µM CHIR-99021, 1 µM SB-431542. On day 7, spheroids were embedded into Matrigel and allowed to continue growing for 7 more days. On day 14, individual spheroids were manually freed from the Matrigel and transferred to a SpinOmega bioreactor spinning at 120 RMP with phase III media: DMEM/F12, 1X N-2 Supplement, 1X B-27 Supplement, 1X Glutamax, 1X MEM Nonessential Amino Acid, 55 µM 2-Mercaptoethanol, 1X Pen/Strep, 2.5 µg/mL insulin. 50 days later, the medium was switched to final differentiation medium: Neurobasal medium, 1X B-27 Supplement, 1X Glutamax, 55 µM 2-Mercaptoethanol, 1X Pen/Strep, 0.2 mM Ascorbic acid, 0.5 mM cAMP, 20 ng/mL brain-derived neurotrophic factor (BDNF), 20 ng/mL glial-derived neurotrophic factor (GDNF). 107-day-old organoids were dissociated into single cells using a 50:50 mixed of Accutase and 0.25% Trypsin with DNase I (1 mg/mL) and plated onto chambered glass slides pretreated with 1% Matrigel (Corning, Inc). Alternatively, organoids were fixed and sectioned for immunostaining (see below). For drug treatment, forebrain organoids (87 Day) were transferred to a 12-well plate on an orbital shaker (120 RPM) and treated with 20 nM KPT276 in final differentiation medium. Medium was changed every other day (Dexoregen, Inc). The organoids were harvested and frozen after 35 days of treatment for sectioning and immunostaining. Each experiment was performed with two batches of organoids per condition to reduce the effect of batch variation. The wildtype cells and K181E mutant have the same genetic background.

### Tissue preparation and immunostaining

Organoids were fixed in 4% paraformaldehyde in PBS for 1 hour at room temperature, washed with PBS, and immersed in 15% sucrose/PBS solution overnight. Subsequently, organoids were embedded in O.C.T. compound (Sakura) in a plastic mold and frozen down in an ultralow freezer. Embedded organoids were sliced onto glass slides using a Cryostat (Leica). Slides were rinsed with PBS, permeabilized with 0.5% Triton-X/PBS solution for 1 hour at room temperature and blocked using 1% donkey serum in 0.1% Tween-20/PBS solution for 1 hour. Primary antibodies, chicken anti-TUJ1 (Abcam, Ab41489, 1:1,000 dilution), rabbit anti-TDP-43 (10782-2-AP, 1/1000 dilution), mouse anti-phospho-TDP-43 (Cosmobio, CAC-TIP-PTD-M01A, 1/500 dilution), were added to the slides and the slides were incubated in a humidified chamber at 4 °C overnight. After several washes in 0.1% Tween-20/PBS solution, DAPI (Sigma) or secondary antibodies, donkey anti-rabbit, anti-mouse, and anti-chicken (Jackson ImmunoResearch), diluted 1:1,000 in blocking solution, were added to the slides. After 1 hour of incubation at room temperature, the slides were washed and mounted in antifade mounting solution (Fisher scientific).

### Image acquisition and processing

Fluorescence confocal z-section images were acquired using a Nikon CSU-W1 SoRa microscope equipped with temperature and CO_2_ control enclosures and a 60x TIRF lens. Randomly selected fields were imaged each with ∼25 slices. Maximum intensity projection views were reconstructed and analyzed by the open-source Image J software (also named Fiji). 3-D and time lapse visualizations were achieved by the Imaris software. Fluorescence intensity was quantified using Fiji. To automatedly count anisosomes, maximum intensity projection views were split into individual channels by Fiji. A consistent default threshold method was applied to each channel. The particle analysis function was used to automatically identify anisosomes or other fluorescence structures for size and intensity measurement.

Statistical analyses were conducted using Excel (for Student’s t-test) or GraphPad Prism versions 8.0, 9.0, 11. P-values were calculated using Student’s t-test in Excel or one-way ANOVA in GraphPad Prism. Curve fitting, including linear and nonlinear models, as well as IC_50_ calculations, were also performed using GraphPad Prism versions 8.0, 9.0 and 11.

### Statistic and reproducibility

All statistical analyses were conducted with GraphPad Prism v10 or Excel. Statistical methods and the number of cells or brain organoid samples (N) are indicated in figure legends or shown in figures as individual data point. Independent repeats (n) are specified in figure legends. No experiment was excluded from the analyses. All experiments were repeated at least twice with individual data point labeled in figures unless specified. For imaging analyses, cells in randomly selected fields were analyzed. The researchers were not blinded. The iPSC cell line was derived from a ADRDs genetic risk-free clone, which was initially obtained from a male. Figures were prepared using ImageJ 1.54f, Imaris 9.9.0, Adobe Photoshop v25.12.1, and Adobe Illustrator 28.7.4.

## Author contributions

YY conceived and designed the study. YY, KC and WZ supervised the study. QZ and KC conducted the chemical and siRNA screens. NC performed post-screen validation and characterization of anisosomes under different conditions. J Z constructed the iPSC knock-in lines and QZ conducted the organoid study. QZ, NC and YY analyzed the data. YY wrote the manuscript. All authors read, edited, and approved the manuscript.

## Corresponding authors

Correspondence to Yihong Ye.

## Acknowledgements

We thank the NIDDK Advanced Imaging Core with data acquisition. We thank D.W. Cleveland (UCSD) and H. Yu (Genentech) for providing the doxycycline-inducible DLD1 TDP-43 2KQ-Clover cell line and the protocol. We thank Y. Xu for technical assistance during revision, K. Bharti and W. Li (NEI), Dr. T. Zhang (NIA) for critical reading of the manuscript. We also thank the NCATS Compound Management Group and the NCATS Automation Group for their assistance in the chemical and genetic screens.

