## Supplementary Figures and legends for "Cellular modifiers of TDP-43 phase transition and cytoplasmic aggregation" (from another version of this preprint: version 2, 2026-04-12)

**Supplemental Materials**

Movie S1 PlaB treatment induces anisosome fusion.

Movie S2 Blocking XPO-1-mediated nuclear export induces anisosome fusion.

**Supplemental Tables**

Table S1 A chemical genetic screen identified compounds that reduce anisosome numbers.

Table S2 A genome-wide siRNA screen identified potential anisosome regulators.

Table S3 Top 110 genes confirmed from a secondary screen as modulators of anisosome dynamics.

**Supplemental Figures**

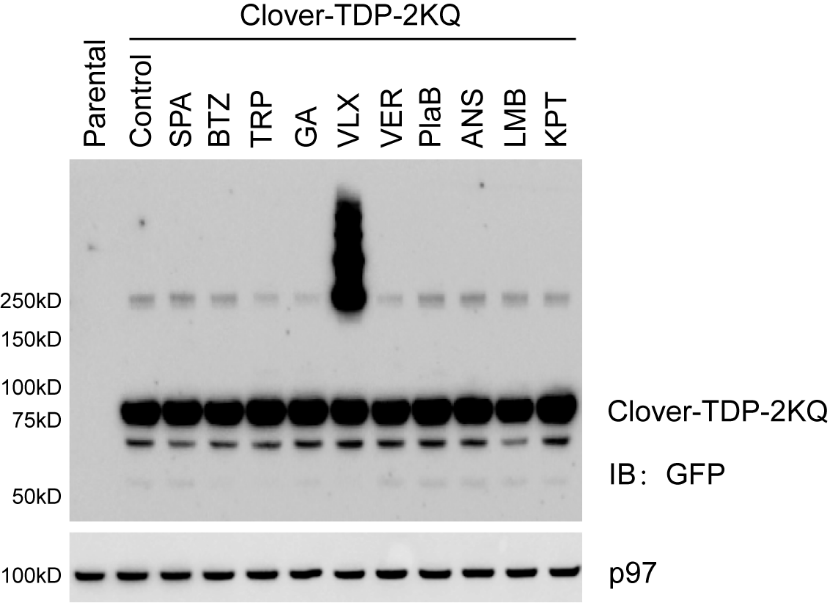

**Figure S1. The impact of different inhibitors on TDP-43 2KQ levels.**

Parental DLD1 or Clover-TDP-43 2KQ cells were exposed to doxycycline for 24 h and then treated with the indicated inhibitors for 6 h. Cell lysates were analyzed by immunoblotting with GFP and p97 (loading control) antibodies. SPA, Spautin-1, 5 μM; BTZ, Bortezomib, 10 nM; TRP, Tripterin, 1 μM; GA, Geldanamycin, 10 μM; VLX, VLX-1570, 0.5 μM; VER, Verdinexor, 20 μM; PlaB, Pladienolide-B, 20 nM; ANS, Anisomysin, 200 nM; LMB, Leptomycin B, 200 nM; KPT, KPT-276, 15 μM.

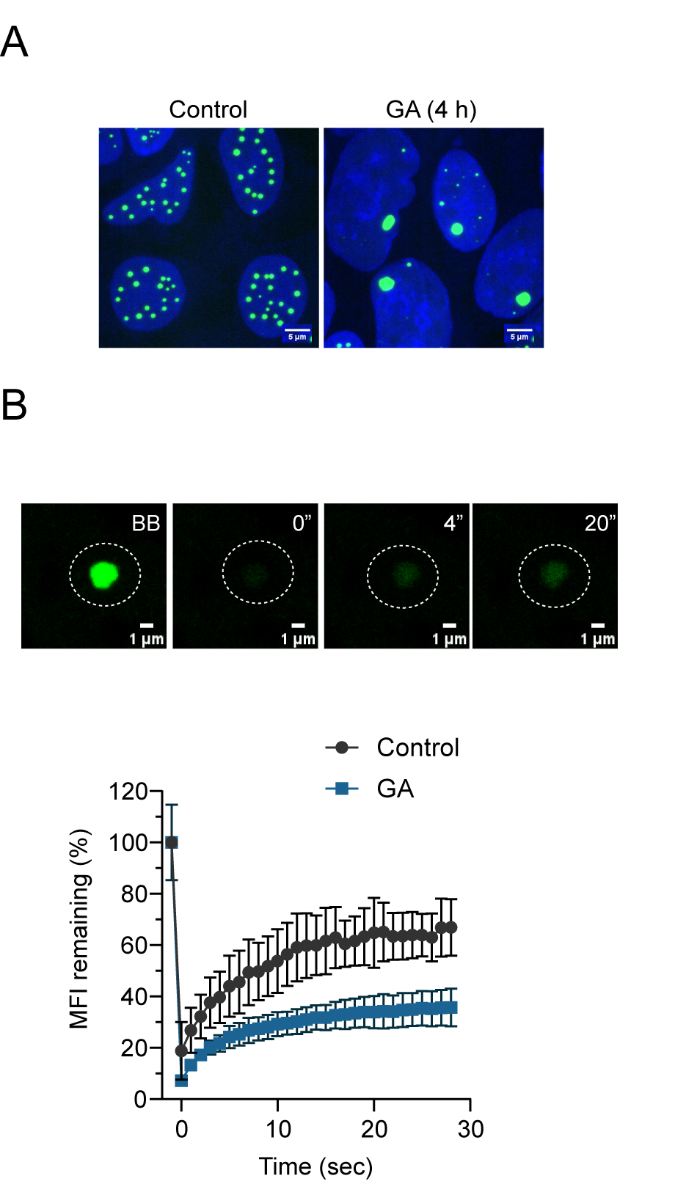

**Figure S2. HSP90 inhibitor Geldanamycin converts TDP-43 anisosome to a gel-like state**

**(A)** DLD1 cells were treated with doxycycline to induce anisosome formation and then treated with Geldanamycin (GA) at 10 μM for 4 h. Cells were stained with Hoechst (Blue) and imaged by a confocal microscope. Scale bar, 5 μm.

**(B)** A TDP-43-bearing anisosome (dashed circle) in a GA-treated cell was photobleached and then imaged. Scale bar, 1 μm. The graph shows the quantification. Error bars, s.e.m. (N= at least 5 anisosomes/condition). MFI, Mean Fluorescence Intensity.

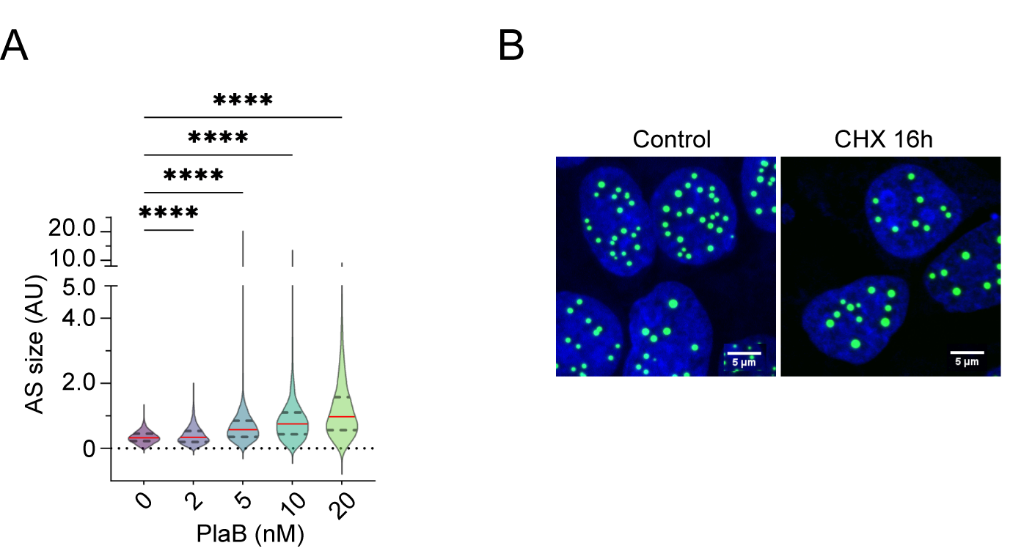

**Figure S3. The impact of splicing inhibitor and translation inhibitor on anisosomes**

**(A)** A violin plot showing the relative size distribution of anisosomes (AS) in DLD1 cells treated with Pladienolide-B at the indicated concentration for 16 h. ****, p<0.0001 by one-way ANOVA. N=3 independent biological repeats.

**(B)** DLD1 cells were treated with doxycycline to induce anisosome formation and then treated with Cycloheximide (CHX) at 20 μg/mL or DMSO as a control for 16 h. Cells were stained with Hoechst (Blue) and imaged by a confocal microscope. Scale bar, 5 μm.

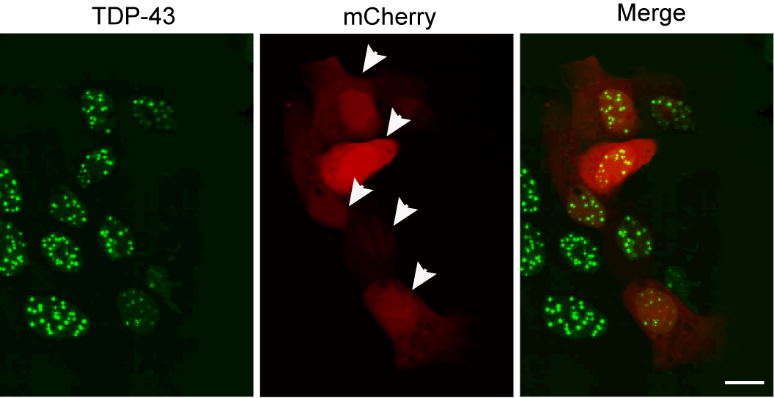

**Figure S4. mCherry overexpression has no effect of TDP-43 phase behavior**

DLD1 cells were transfected with a mCherry-expressing plasmid and then induced for TDP-43 expression for 24 h before imaging. The arrowheads indicate mCherry-positive cells that have only nucleus-localized anisosomes. Scale bar, 10 μm.
